# Is a Picture Worth 1,000 SNPs? Effects of User-Submitted Photographs on Ancestry Estimates from Direct-to-Consumer Canine Genetic Tests

**DOI:** 10.1101/2023.06.28.546898

**Authors:** Halie M. Rando, Kiley Graim, Greg Hampikian, Casey S. Greene

## Abstract

**Objective:** To evaluate whether the breed ancestry predictions of direct-to-consumer (DTC) genetic tests for dogs are influenced by the user-provided photograph.

**Animals:** Twelve pet dogs considered purebred (i.e., registered with a breed organization) representing twelve different breeds.

**Methods:** Six buccal swabs per dog were collected by the owners and submitted to six DTC genetic testing companies. The experimenters registered each sample with the company. For half of the dogs, the registration included a photograph of the DNA donor. For the other half of the dogs, photographs were swapped between dogs. Analysis of the DNA and breed ancestry prediction was conducted by each company. Each company’s breed predictions were evaluated to assess whether the condition (i.e., matching versus shuffled photograph) affected the odds of identifying the DNA donor’s registered breed. A convolutional neural network was also used to predict breed based solely on the photograph as a positive control.

**Results:** Five of the six tests always produced results that included the registered breed. One test and the convolutional neural network were unlikely to identify the registered breed and frequently returned results that included the breed in the photograph. This result suggests that one test on the market is relying on the photograph more than the DNA sample. Additionally, differences in the predictions made across all tests underscore the challenge of identifying breed ancestry, even in purebred dogs.

**Clinical Relevance:** Veterinarians are likely to encounter patients who have conducted DTC genetic testing and may find themselves in the position of explaining genetic test results that they did not order. This systematic comparison of tests on the market provides context for interpreting unexpected results from consumer-grade DTC genetic testing kits.

## Introduction

Clinical genetic testing is an invaluable tool for human and veterinary medicine^1,2^. Though clinical veterinary medicine has emerged relatively recently, the study of genetic disease in animal models has a long history, and commercial genetic testing has been used to guide agricultural breeding practices for over 20 years^3^. Commercial testing for genetic risk factors has been common for dogs^4^ and cats^5^ for over a decade, and animal genetic testing is a booming industry that is expected to grow over the next five years^6^. Increasingly, the decision to conduct genetic testing of both humans and companion animals occurs outside of the clinic.

Over the past 30 years, direct-to-consumer (DTC) genetic testing of humans has grown into an established industry^7^. These tests allow consumers to collect a specimen (usually a buccal swab) at home and ship it to the company to be processed, sequenced, and analyzed. The company then returns a report about the individual’s ancestry and, in some cases, the likelihood of possessing genetic traits (e.g., hair color) or risk of developing genetic diseases. Recently, a parallel industry of DTC genetic profiling of companion animals, primarily dogs, has also emerged. The dog equivalent of these ancestry estimates is the identification of the breed make-up of an individual pet. Dogs were the first domesticated animal^8^, and dog husbandry has a rich history^9,10^. As a result of dog breeding practices, modern dog breeds are more differentiated than subpopulations of many other species, and individual breeds are associated with distinct genetic signatures^11^, allowing the breed ancestry of a particular dog to be estimated based on the presence of genetic variants associated with specific breeds. Additionally, the genetic etiology of many canine diseases is known (e.g.,^12^), allowing DTC genetic testing companies to predict individual dogs’ risk of developing various diseases.

For companion animal DTC genetic tests, the consumer collects a buccal swab from a pet that they ship back to the company, which sequences and analyzes the animal’s DNA. Most DTC genetic tests assay single-nucleotide polymorphisms using microarray-based profiling, though some use other genotyping techniques (see Table 1). In all cases, the stated goal is to identify genetic variants in an individual dog that can be compared against a panel of genetic variants sampled from a variety of breeds. This approach allows for classification of the individual’s genetic variants to one or more breeds, which can then be deconvolved into measured sources of genetic ancestry. The genotypes can additionally be analyzed for association with certain traits or diseases (usually for an extra fee). Therefore, these analyses depend on three variables: first, the density of markers analyzed; second, the set of breeds included in the reference panel; and third, and diversity of individuals from each breed available within the panel. Typically, however, the specific approach used is proprietary, and the consumer receives only a report outlining their dog’s estimated breed ancestry and/or the presence of known genetic risk variants. Therefore, the specifics of how these estimates are made and their accuracy remains largely unknown.

**Table 1:** Comparison of six DTC dog genetic testing services. At the time of study design (end of 2021), six DTC dog genetic tests were commercially available. All six were included in this study. Each test uses a distinct combination of genotyping technologies, breed classification algorithms, and reference panels. While this study focuses on the methodologies, a detailed breakdown of consumer-oriented similarities and differences among several of these tests is available elsewhere^18^.

| Test/Company Name | Price of Base Breed Test | Markers Used | Reference Panel | Ancestry Assignment Algorithm |
| --- | --- | --- | --- | --- |
| Embark Breed ID DNA Test<sup>19</sup> | \$129-\$199 | 200,000+ SNPs, custom chip<sup>20,21</sup> including all markers (173k) on the Illumina CanineHD platform<sup>22</sup> | 350+ breeds<sup>23</sup> | Not specified |
| Wisdom Panel Breed Discovery<sup>24</sup> | \$80-\$160 | SNPs, number not specified, custom-designed Illumina Infinium XT microarray<sup>12</sup> | 350+ breeds, 21,000+ samples<sup>12,25</sup> | BCSYS<sup>12</sup> |
| Orivet GenoPet<sup>26</sup> | \$95-160 | SNPs, number not specified<sup>27</sup> | 15,000 samples from 350+ breeds<sup>28</sup> | Not specified |
| Darwin's Ark Explorer Level Donation<sup>29</sup> | \$149 donation | Call 9M SNPs from whole-genome sequencing, use 688K for breed classification<sup>30</sup> | 101 breeds<sup>31</sup> | SupportMix<sup>32</sup> |
| DNA My Dog Essential Breed ID Test PLUS Wolf - Canid/Hybrid Test<sup>33</sup> | \$80-\$130 | Copy-number variation, number of markers not specified<sup>34</sup> | 350+ breeds<sup>35</sup> | Not specified |
| Accu-Metrics/Via-Pet Canine Breed Determination<sup>36</sup> | \$59 | Not specified | 340 breeds<sup>36</sup> | Not specified |

This lack of transparency raises questions, especially in cases where different tests produce different results. Crowd-sourced DTC dog genetic testing results, available through the “DoggyDNA” subreddit^13^, suggest that some dogs receive wildly different breed assignments from different companies, beyond what would be expected based on the known methodological differences. One potential source of variation could be that all companies either allow or require the user to upload a photograph of the dog, despite the tests being advertised as based on genetics only. If the photographic information is incorporated into the breed identification pipeline, it could bias the results. Unfortunately, because most of the dogs whose results are uploaded to the DoggyDNA subreddit are mixed-breed, it is impossible to evaluate the ground truth using these crowd-sourced results to compare different companies’ genetic ancestry estimates. Therefore, we sought to assess whether the photograph uploaded with a purebred dog’s DNA sample influenced estimates of their breed ancestry.

We evaluated the six DTC tests on the market at the end of 2021 (when our study began). We recruited twelve purebred dogs and submitted their buccal DNA samples along with a photograph to each company. For half the tests, the DNA sample was submitted with a photograph of the DNA donor. For the other half, photos were between dogs so that each DNA sample was submitted with a photograph of a different purebred dog. To evaluate whether the photograph affected ancestry estimation, we checked whether predicted breeds matched the donor’s registered breed. We also evaluated samples using a photograph-only baseline that used a deep neural network trained to identify dog breeds^14^. This design allows us to provide a formal comparison of differences in breed ancestry estimation across different platforms.

## Methods

Dogs were recruited by word of mouth in several locations between July and November 2022. Breed organization registration of each dog was confirmed by manually viewing the dog’s breed organization paperwork or locating the dog in the American Kennel Club (AKC) database. Recruitment was structured to ensure full coverage across dog breed clades^15^. Each dog was randomized to either the control or treatment group, corresponding to the dog’s own photo versus a randomized photo. Dogs in the treatment group were paired based on their enrollment order, and the photographs were swapped across pairs. Assignments were evaluated to ensure that dogs within a breed clade (per Parker et al.^15^) were not paired. All enrolled dogs came from different households, avoiding the possibility of cross-contamination of dogs within the study.

The six DTC genetic tests (Table 1) were purchased directly from each company’s website. Kits were registered according to the instructions provided, using the dog’s own photograph (control) or the photograph of a paired dog (treatment). When the information was requested, dogs were listed as mixed breed. A set of kits was then delivered to each dog owner, who was instructed to collect buccal swabs for each according to manufacturer instructions and to mail the samples using the company-provided packaging. Genetic testing was conducted by each DTC genetic testing company according to their own protocols. The breed determination results were returned to the authors, who retrieved the results for each dog and recorded all results for the determined breeds and the corresponding percentage composition in a spreadsheet. Breed results were harmonized across studies using the Vertebrate Breed Ontology version 20230601^16^. If a breed assignment corresponded to a variety, rather than a breed, per the Federation Cynologique Internationale (FCI)^17^, the corresponding breed (i.e., the parent term) was used instead. Qualitative assignments, such as ranges of percentages, were converted to numerical values for the purposes of data visualization (Supplementary Table 2).

As a positive control for the potential photographic test bias, we predicted the breed of each sample/photograph pair using a pre-trained convolutional neural network. Dog breed classification is a classic problem in computer vision, and models trained on images from a large compendium (ImageNet) have been evaluated for their performance in classifying photographs of purebred dogs. The model NASNet shows particularly robust performance^37–39^. Therefore, we used NASNet predictions to complement the DTC genetic testing results by predicting breeds based exclusively on photographs. We loaded the NASNet model **nasnetalarge** from the Python package PyTorch Image Models (timm version 0.6.12)^40^ and used it to it to classify each of the dog photos, adapting code from a tutorial^41^. The classifier estimates many categories with a very long tail, and so only hits with a score >1% were included in the analysis.

To evaluate whether swapping the photographs affected breed prediction for each DTC test, the proportion of results matching the registered breed across the two conditions was compared by calculating the odds ratio. In cases where there were either zero matching or zero non-matching predictions within a condition (i.e., one or more zero cells), the Haldane-Anscombe correction^42^ was applied per^43,44^. We evaluated independence between the treatment and control conditions using a one-tailed hypergeometric test. Rejection of the null would indicate a potential relationship between the treatment (photograph matching status) and outcome. Both statistical tests were conducted in R using the package vcd^45,46^.

Based on qualitative patterns observed in the results, the decision was made *post hoc* to compare the phylogenetic clade^15^ of each predicted breed against the registered breeds of the DNA donor and photographed dog. Each breed was mapped onto a clade, as identified by Parker et al.^15^ or, for breeds that were not included in that analysis, based on its closest genetic relationship to breeds assigned to clades. In the treatment condition, each predicted breed’s clade was compared to the clades of the DNA donor versus the dog whose photograph was provided to the testing company. In the control condition, each dog was paired with another dog (based on the order in which they were recruited) to provide a random baseline. The breed predictions were then evaluated to determine whether their clade matched the clade of the photographed or paired dog in the treatment and control conditions, respectively. The goal of this analysis was to evaluate whether predicted breeds that matched neither the registered breed of the DNA donor nor the photograph were closely related to but not identical to either dog breed. The odds ratio and hypergeometric tests were applied using the vcd package in R. Additionally, the results at the level of breed clade were plotted in R using the ternary package^47^. Qualitative results (e.g., range-format percentages) were converted to numerical values for the purposes of this visualization (Supplementary Table 2).

## Results

Twelve purebred dogs were recruited from households around the United States to have DNA samples collected by their owners (Table 2). Most results were returned from the DTC genetic testing companies between August 2021 and February 2023. Some results from Orivet were delayed due to an issue with the user interface and were returned in June 2023. Results were not received for four samples. One, sent to Darwin’s Ark, was lost between leaving the dog owner’s house and being processed, and the company was responsive in providing updates and offering to replace it, although based on timing it was ultimately dropped from the study. The remaining three lost samples were sent to Accu-Metrics, and as of June 2023, no status update could be obtained from the company.

**Table 2:** Background information on the twelve canine participants. All dogs were registered with a breed organization (AKC = American Kennel Club; UKC = United Kennel Club; CKC = Continental Kennel Club). Breed clade was assigned based on phylogeny^*15*^ as opposed to the breed groups used by breed organizations such as the AKC.

| Breed | VBO Identifier | Breed Clade | Registry Organization | Condition | Photograph |
| --- | --- | --- | --- | --- | --- |
| Beagle | VBO:0200131 | Scent Hound | UKC | Control | Self |
| Border Terrier | VBO:0200194 | Terrier | AKC | Control | Self |
| Golden Retriever | VBO:0200610 | Retriever | AKC | Control | Self |
| Pomeranian | VBO:0201043 | Small Spitz | AKC | Control | Self |
| Shetland Sheepdog | VBO:0201217 | UK Rural | AKC | Control | Self |
| Shih Tzu | VBO:0201223 | Asian Toy | CKC | Control | Self |
| Brittany | VBO:0200238 | Pointer/Setter | AKC | Treatment | Chinese Crested |
| Chinese Crested | VBO:0200345 | American Toy | AKC | Treatment | Brittany |
| German Shorthaired Pointer | VBO:0200583 | Pointer/Setter | AKC | Treatment | Italian Greyhound |
| Italian Greyhound | VBO:0200713 | UK Rural | AKC | Treatment | German Shorthaired Pointer |
| Bulldog | VBO:0200258 | European Mastiff | AKC | Treatment | Labrador Retriever |
| Labrador Retriever | VBO:0200800 | Retriever | AKC | Treatment | Bulldog |

For all dogs, the registered breed was always the majority estimate, and at least one DTC test returned a result 100% consistent with the registered breed (Figure 1; Supplementary Table 4). In most cases, at least one test’s breed prediction did not match the registered breed. Most tests returned at least one result that did not match the dog’s breed organization registration. Our analysis focused on evaluating whether the reported ancestry deviated from the expected ancestry, which in purebreds is 100% of a single breed, so minor differences in how results were reported by different companies did not affect our analyses.

**Figure 1:**
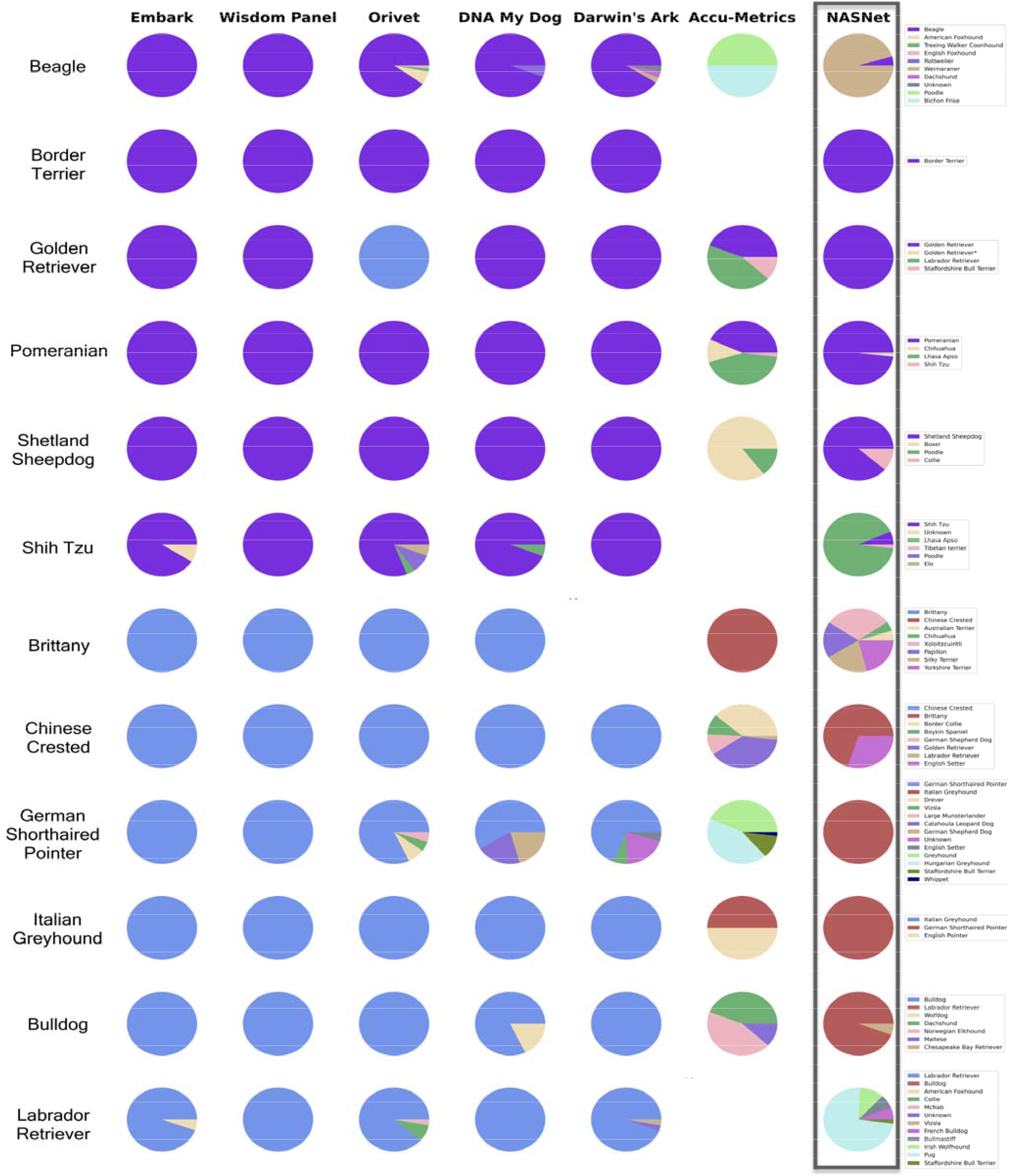
Breed ancestry predictions from six DTC dog genetic testing companies as well as a pretrained image classifier. For the control condition, the breeds of the donor and photograph match, so the registered breed is depicted as purple. In the treatment condition, these are separated into blue (DNA donor) and red (photographed breed). All other breeds are represented in other colors as identified in the legend.

All but 4 of the breed estimates mapped unambiguously to the VBO (Supplementary Table 1). In one case, a result returned by NASNet (“Lhasa”) is not a known dog breed. “Lhasa Apso” was assumed to be the intended breed. In the other three cases, two or more potential VBO terms could correspond to the breed result. In two cases, VBO developers were consulted to select the best mapping. The last case was Catahoula Leopard Dog, a dog breed with a complex history that could correspond to three VBO entries. However, this result was returned only by one test for one dog, negating the need for standardization, and we simply selected the closest VBO term to the exact phrasing used in the breed result. The translation of qualitative to quantitative results is provided in Supplementary Table 2, and the standardization of NASNet’s predictions are shown in Supplementary Table 3.

Another difference in the way results were reported across tests was that Darwin’s Ark provided a two-component breed prediction. As Darwin’s Ark evaluates ancestry and genetic diversity separately, they provided not only the most genetically similar breeds but also an explicit prediction of whether a dog was likely to be purebred. We used these determinations of breed, rather than the ancestry percentages (Figure 1), to assess whether Darwin Ark’s result matched the registered breed (Table 3). Of the 11 samples for which results were returned, Darwin’s Ark successfully discerned that all donors were purebred, even when some genetic variants were assigned to other breeds (Table 3; Figure 1).

**Table 3:**
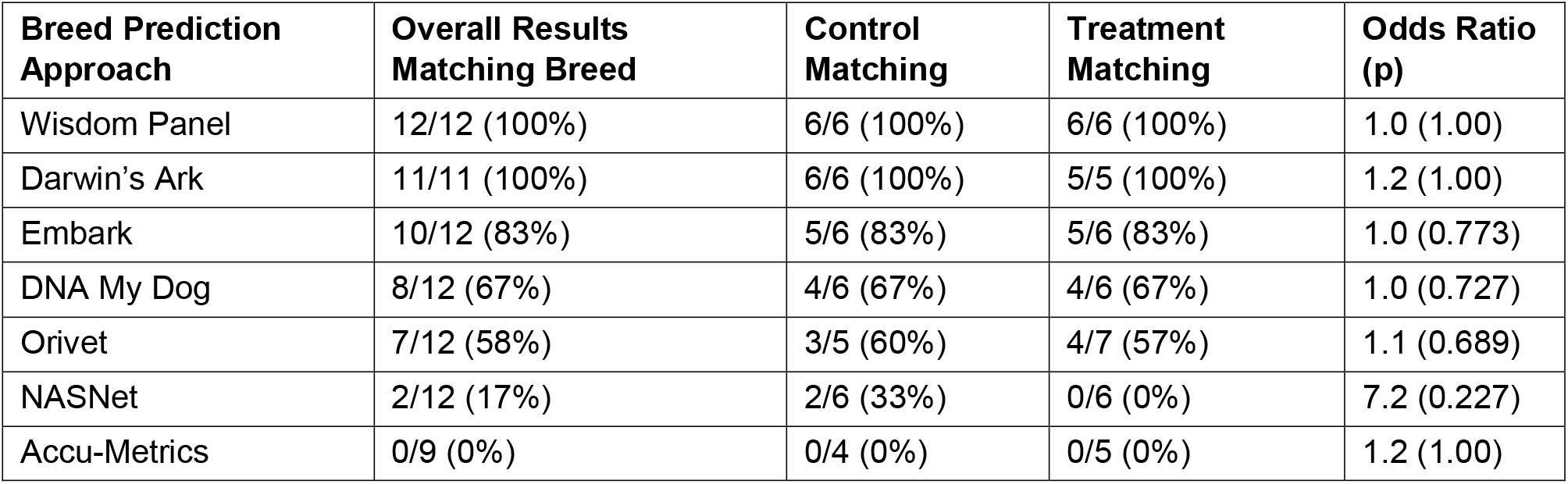
Comparison of six DTC genetic tests and one pretrained image classifier (NASNet) to the registered breed of each dog. The overall concordance with breed registration and the breakdown between the two conditions are reported. The Haldane-Anscombe correction was applied when one or more cells were zero. P-values were calculated using a one-tailed hypergeometric test. Note: for Orivet, the conditions were not balanced because, due to experimenter error, one control subject (Golden Retriever) was submitted with two photographs (the photograph of the DNA donor and of the Pomeranian). Also, due to sample loss, only 11 dogs were tested using Darwin’s Ark and 9 for Accu-Metrics versus 12 for the other DTC tests.

We note that the primary outcome selected *a priori* (Table 3) did not capture apparent qualitative differences (Figure 1). This discrepancy was driven by the fact that, in several cases, the breeds predicted by a test matched neither the DNA nor the photograph. This pattern was particularly apparent for the image classifier and one DTC genetic test (ViaPet by Accu-Metrics). In the case of the neural network, this was likely influenced by the limited number of breeds in the training data, and thus category labels, in ImageNet^48^. For example, ImageNet is not trained to identify Chinese Crested and Bulldog breeds. In the case of the genetic test, the results may have been influenced by the fact that the test predicted the registered breed zero times across both the control and treatment conditions (Table 3). Re-evaluating the results at the level of breed clades (Figure 2) revealed that, in both cases, the predicted breeds were more similar to the photographs provided than to the DNA sample analyzed. While the effects did not meet the level of statistical significance (Supplementary Table 5), the odds ratios indicate that the treatment condition had a similar effect on the ability of NASNet and Accu-Metrics to predict the breed of the DNA donor (odds ratio of 13 versus 11). Given that NASNet assigned breeds based solely on photograph, it was expected to incorrectly predict the treatment samples (which had shuffled photos). The fact that a similar effect was observed for predictions made by Accu-Metrics suggests that the photograph was also influential in the analysis made by this DTC dog genetic test.

**Figure 2:**
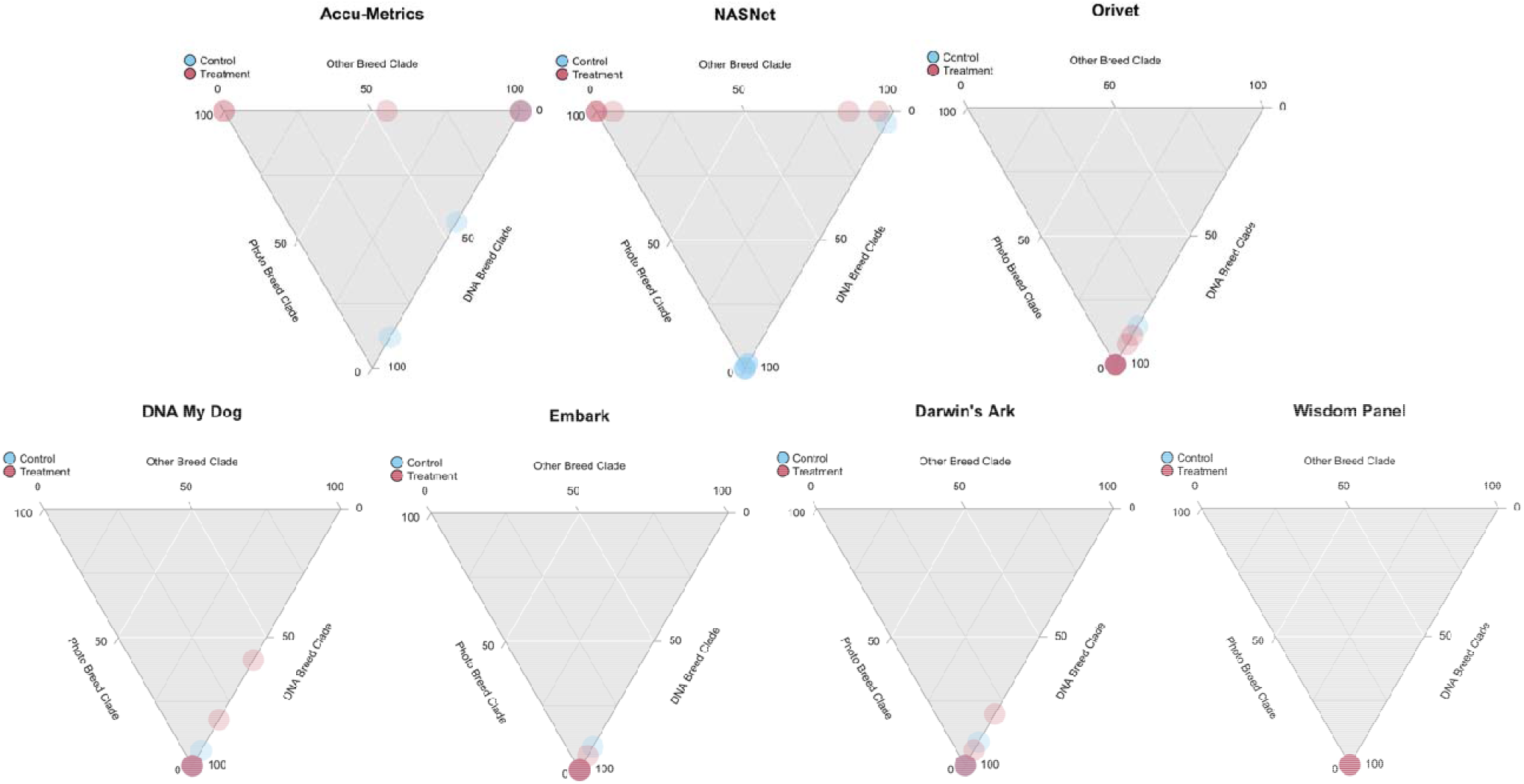
Ternary plots identifying the breed clades of results predicted by each test. Each dog is represented by a circle in blue (control) or red (treatment). The position of the circle relative to the three axes indicates the percentage of results belonging to the breed clade of the DNA donor, photographed dog, or other clades. A result that is 100% concordant with the DNA donor’s registered breed will be placed at the apex of the triangle. A result that is 100% concordant with the photographed dog’s breed (in the treatment condition only) will be located at the top left. Offset from these positions can be interpreted by looking at the position relative to the other two vertices on a 0 to 100 scale. Darker colors indicate multiple circles mapping to the same position.

## Discussion

This study applied an experimental paradigm to assess the state of DTC genetic testing for dogs and revealed qualitative and quantitative differences in the breed ancestry estimates of six providers. By modulating whether DNA was submitted with a matched or mismatched photograph, we evaluated whether this non-genetic information influenced breed predictions. Using a pre-trained convolutional neural network, NASNet, allowed us to compare the predictions made by DTC genetic tests to those made from the photograph alone. Ultimately, this analysis demonstrated a low rate of consensus across tests in estimating the breed makeup, even of registered purebred dogs. The results suggest that at least one company is providing results influenced by the photograph provided.

Both NASNet and Accu-Metrics were unlikely to identify the registered breed as 100% of the ancestry, regardless of condition. When breed ancestry estimates were evaluated at the clade level, there was a significant effect of the photograph on the predictions made by NASNet and Accu-Metrics; the clades of the predicted breeds matched that of the dog in the photograph instead of the DNA donor (Figure 2). For NASNet, which is an image recognition tool, this result was expected; however, Accu-Metrics collects a DNA sample and even rejected one submission for failing quality control. The lack of a relationship between the company’s breed predictions and the provided DNA samples suggests that Accu-Metrics used the photograph, rather than the DNA, to make its predictions.

In overall performance, Accu-Metrics was less successful in predicting a dog’s breed than NASNet, with an overall rate of 0% versus 17% (Table 2). NASNet may have had even higher performance if more breeds were included in its training dataset, given that it did not have any point of reference for some of the breeds examined in this study (e.g., Chinese Crested). The Chinese Crested photograph was, incidentally, the only photograph that Accu-Metrics identified as 100% matching the registered breed of the photographed dog (Supplementary Table 4); in this case, however, the photograph of the Chinese Crested was submitted along with DNA from a Brittany spaniel. Therefore, this study suggests that Accu-Metrics is not providing accurate information about genetic ancestry, although the product has been promoted by companies such as the pet pharmacy PetMeds^49^.

Among the remaining five DTC testing services, the rate of predicting the registered breed of the DNA donor was very high, regardless of the condition. This result suggests that these companies are indeed analyzing the DNA as the basis for their breed ancestry predictions. Additionally, because the statistical analysis did not support an effect of condition on the odds that the photographed breed (or its close relatives) was included in the predictions, the results suggest that these tests are not biased by the photographs.

While it is interesting to explore differences in the results produced by different tests, it is important to note that this study was not designed to evaluate the accuracy of the tests’ genetic ancestry predictions. This point is especially relevant to tests with extremely high performance. While Wisdom Panel returned a prediction that matched the DNA donor’s registered breed 100% of the time, this does not mean it is necessarily more accurate than Embark or Orivet. Differences among tests, such as the number of markers analyzed (Table 1) are expected to contribute to differences in test resolution and breed prediction capabilities. Additionally, the specific breeds and specific dogs included in a company’s reference panel can have a strong effect on breed assignment.

Among the twelve purebred dogs tested, some received more consistent predictions than others. For more than half the dogs, there was consensus among five DTC genetic tests (excluding Accu-Metrics) about their breed (Figure 1). Among the others, though the registered breed was always the major ancestry estimate from each of the five DTC tests, they often differed in what other breeds were predicted to contribute to ancestry. Pedigrees are not infallible, and therefore mistaken parentage could be a source for some of this variation. However, it also underscores the complexity of the question of genetic ancestry versus a socially defined population, i.e., breed. Therefore, even with the ground truth provided by using registered dogs, some sources of genetic variation are difficult to determine.

One notable example comes from the German Shorthaired Pointer (GSP), which belongs to the Pointer/Setter clade^15^. In this study, the GSP sample was submitted with a photograph of an Italian Greyhound, which belongs to the UK Rural clade^15^. Embark and Wisdom Panel each identified this dog as 100% GSP (Figure 1). Orivet and Darwin’s Ark both identified GSP as the dog’s primary ancestor (82% and 69%, respectively), but also included other Pointer/Setter breeds such as the Vizsla^50^ in their ancestry estimates. In addition to the GSP, Vizsla, and an additional 20% of the dog’s genetic variation that could not be assigned to a breed, Darwin’s Ark also predicted Drever ancestry; the Drever is a Swedish scent hound related to the Dachshund^50–52^, introducing another breed clade. Alternatively, DNA My Dog identified 59% of the dog’s ancestry as GSP and the remainder as German Shepherd Dog and Catahoula Leopard Dog, both of which are North American breeds^15,53^. Thus, while these three tests all predicted that 20-40% of the genome was indicative of non-GSP ancestry, there was substantial variation in the assignment of this DNA across tests. While it is not possible to extrapolate why the three tests arrived at different predictions, the fact that both Wisdom Panel and Embark, which are the most popular and comprehensive DTC canine genetic tests, identified the dog’s ancestry as 100% GSP suggests that the dog carries some less-common GSP genetic variation that is not present in the comparison panels used by the smaller companies. Less-common variants in this individual dog, combined with differences in the sample size of the reference panel, could explain why both Darwin’s Ark and Orivet assigned some DNA to non-GSP breeds within the Pointer/Setter clade. However, it would explain neither the Drever result from Darwin’s Ark nor the New World breeds assigned by DNA My Dog.

Similar narratives also arose from the ancestry predictions for two other dogs. In the case of the Beagle, Embark and Wisdom Panel agreed on 100% Beagle ancestry, while Orivet, DNA My Dog, and Darwin’s Ark all reported >90% Beagle ancestry. The sources of non-Beagle ancestry were scent hounds and/or pointer/setters or drovers. In the case of the Shih Tzu, there was convergence among tests on potential Asian-Toy or Poodle ancestry that accounted for 5-20% of genetic ancestry (Supplementary Table 4).

There was also a lack of consensus among DTC tests for the Labrador Retriever. Wisdom Panel and DNA My Dog were the only tests to identify her as 100% Labrador Retriever. Embark predicted that she carries 5.5% American Foxhound DNA, suggesting a Foxhound ancestor approximately five generations ago. This individual dog’s pedigree indicates that six of her sixteen great-grandparents were enrolled in the AKC’s Voluntary DNA Profile Program, which allows breeders to confirm their dogs’ parentage. Therefore, the foxhound admixture is unlikely to come from these six ancestors. However, among the remaining ten great-grandparents, it is possible that one had mistaken parentage and an American Foxhound parent, especially since Embark’s chromosome chart for this dog suggested inheritance of Foxhound ancestry from only one parent. The other high-resolution test, Darwin’s Ark, estimated a similar amount (5.9%) of non-Labrador ancestry for this dog. However, Darwin’s Ark does not currently include the American Foxhound in its reference panel, and instead assigned this non-Labrador ancestry to Viszla (3.0%) and unknown (2.9%). Notably, although Orivet includes the American Foxhound in its reference panel, it identified 11.3% of this dog’s ancestry as Collie (8.4%) and McNab (2.9%), but not American Foxhound. The Collie is a UK Rural breed, while the McNab Shepherd is an American creation bred from Collies and Basque shepherd dogs of unknown ancestry^54^. Thus, once again, no simple story can be discerned by comparing the results across different tests. This Labrador may have one great-grandparent with non-Labrador ancestry, but no consensus arose about what this ancestry could be. In contrast to the GSP, however, none of the tests assigned the ancestry to a breed closely related to the Labrador, aligning more with historical admixture than rare breed-specific genetic variants. Here, it is worth noting that Embark and Darwin’s Ark conduct very high resolution analyses due to their marker density (Table 1), providing a possible explanation for why they would detect a ∼5% non-Labrador ancestry when tests like Wisdom Panel did not. It’s, therefore, possible that this result is an illustration of ascertainment bias (a type of sampling bias), where lower-resolution tests either failed to detect any non-Labrador variants or overestimated the non-Labrador contribution. Unfortunately, because the number of SNPs used by several tests (including Orivet) is not publicly available, it is impossible to know definitively whether this is the case.

As a third and final example of disagreement in breed estimation, DNA My Dog was alone in reporting that the purebred Bulldog had 10-20% wolf ancestry. This result would be consistent with a full wolf ancestor three generations ago. While not all tests evaluate the genome for non-dog contributions, Embark, Wisdom Panel, Orivet, and DNA My Dog all do. Therefore, it would be surprising for all the other tests to miss such a high percentage of wolf ancestry. One possible explanation is that DNA My Dog uses copy number variation, whereas the other tests use SNPs (Table 1). Different genetic tools tell different stories. However, a recent media report indicated that DNA My Dog failed to identify the species of a sample, assigning dog breeds to human DNA^55^. While our analysis provides evidence that the company did sequence the DNA samples we submitted, it is possible that other issues (e.g., contamination) could cause issues in species identification and breed prediction.

This analysis demonstrates the complexity of the DTC dog genetic testing market. In one case, the results we received were independent of the DNA submitted. Among the five other tests, the results clearly demonstrate that methodological literacy is essential for interpreting DTC test results. Factors such as the DNA marker type and SNP panel density analyzed, the size and composition of the breed reference panel, and the assignment algorithm can change how variants in the DNA are identified, interpreted, and assigned to breeds and diseases. Unfortunately, the average consumer is unlikely to have the necessary information or training to critically evaluate such aspects when selecting a DTC genetic test^56^.

Lack of public literacy related to genetic testing and transparency in DTC test methodologies is especially concerning in this context, because dog breed ancestry can have social and economic consequences. For example, many home insurance companies refuse to cover certain dog breeds such as Pitbulls (e.g., American Staffordshire Terriers) and Rottweilers. In our current sample of twelve purebred dogs (none of which are banned by more than 5% of insurance companies^57^), four dogs were identified by at least one test as having ancestry banned by more than 50% of companies. DNA My Dog identified the Bulldog as a wolf hybrid and the Beagle as part Rottweiler. Accu-Metrics identified both the GSP and Golden Retriever as part American Staffordshire Terrier. None of these ancestry predictions were supported by any other DTC tests. Therefore, dog owners could face severe financial repercussions in terms of home insurance and even housing rental eligibility if the results of a DTC test are viewed as definitive^58^ or if DTC canine genetic testing are adopted for housing and insurance purposes. Given DTC genetic services are being used by some rental companies for pet waste identification^59^, it is not difficult to envision landlords conducting or requiring breed testing as well^60,61^. Additionally, it is legal for companies to decline to insure or to raise premiums for dog owners who they believe own a restricted breed^62^. While insurance companies do not typically require DNA tests, many sources advise renters and homeowners to submit their dog’s DNA results if they believe they were erroneously rejected for coverage because of their dog^58,63–65^. Notably, several of these websites suggest that canine DNA tests can be purchased for around $50. Currently, DTC dog genetic test results cost on average $99, and Accu-Metrics is the only test available for less than $80. While breed discrimination in renting and insurance is controversial^62^, including inaccurate breed ancestry estimates will only serve to further muddy an already complicated issue.

It is also imperative to consider the context of the other services that many DTC genetic tests provide: health predictions. Despite being priced similarly to human DTC test kits, DTC canine genetic testing is not held to the same regulatory standards as human testing in the United States. Some services (Embark, Wisdom Panel, and Orivet) provide information about specific risk variants that a dog does or does not possess. DNA My Dog and Accu-Metrics, on the other hand, provide general information about health risks based on the breeds detected. The potential for such information to be misunderstood and misapplied has been widely discussed as a general risk of DTC genetic testing^66^. The human DTC market underwent a major shift in the mid-2010s when the FDA began regulating the health results they are allowed to provide to consumers^66,67^. Genetic information can contribute to pet owners’ decisions regarding their pet’s health, and when this information is incorrect or is interpreted incorrectly, it can lead to tragedy^68^. The team of scientists leading Darwin’s Ark has been outspoken about this potential for misuse^68^, and Darwin’s Ark does not provide health predictions because of a lack of rigorous standards for these estimations^30^. The extent to which health predictions differ among tests would also be an interesting question for future studies of the industry. However, the results from this study make it clear that improved regulation is essential to ensure that consumers are receiving accurate information from DTC canine genetic testing companies.

For more than a decade, concerns have been raised about the potential pitfalls of human DTC genetic testing, especially when combined with limited regulatory oversight. Now, veterinary medicine faces several related issues. Our systematic comparison of DTC genetic tests for dogs suggests that it is imperative that consumers approach DTC test results with care. Veterinarians are likely to be placed in a position where they need to educate pet owners about genetic tests that the veterinarians did not order or recommend. Many factors influence the results, potentially resulting in surprising predictions. While we did not analyze health or trait predictions, this study shows that ancestry predictions were often at odds with the dogs’ pedigree registration, even for AKC-registered dogs (10 out of the 12 participants). For one test, the DNA sample submitted had little bearing on the results, whereas the photograph heavily influenced breed predictions. Our findings suggest that DTC genetic test results should be interpreted very cautiously. As DTC testing expands to make predictions about pet health (from dietary needs^69^ to age^70^), veterinarians may face increased calls to educate owners about the limits of genetic testing.

## Supporting information

Supplementary Tables 1-5 (as sheets)

## References

1. Patterson DF. Companion Animal Medicine in the Age of Medical Genetics. Journal of Veterinary Internal Medicine 2000;14:1–9. Available at: https://doi.org/dntg9j.

2. Scott S, Abul-Husn N, Owusu Obeng A, et al. Implementation and utilization of genetic testing in personalized medicine. PGPM 2014:227. Available at: https://doi.org/gr9hjb.

3. Dekkers JCM, Hospital F. The use of molecular genetics in the improvement of agricultural populations. Nat Rev Genet 2002;3:22–32. Available at: https://doi.org/bsq7pk.

4. Bell JS. Researcher responsibilities and genetic counseling for pure-bred dog populations. The Veterinary Journal 2011;189:234–235. Available at: https://doi.org/dr4ht3.

5. Lyons LA. Genetic testing in domestic cats. Molecular and Cellular Probes 2012;26:224–230. Available at: https://doi.org/f4gjgw.

6. Global Market Insights. Animal Genetics Market revenue to cross USD 6.4 Bn by 2027. GlobeNewswire News Room 2021. Available at: https://www.globenewswire.com/news-release/2021/03/08/2188474/0/en/Animal-Genetics-Market-revenue-to-cross-USD-6-4-Bn-by-2027-Global-Market-Insights-Inc.html. Accessed June 13,2023.

7. Hogarth S, Saukko P. A market in the making: the past, present and future of direct-to-consumer genomics. New Genetics and Society 2017;36:197–208. Available at: https://doi.org/gr9hh9.

8. Clutton-Brock J. The process of domestication. Mammal Review 1992;22:79–85. Available at: https://doi.org/b29ssc.

9. Galibert F, Quignon P, Hitte C, et al. Toward understanding dog evolutionary and domestication history. Comptes Rendus Biologies 2011;334:190–196. Available at: https://doi.org/b7dmx4.

10. Jung C, Pörtl D. How old are (Pet) Dog Breeds? Pet Behav Sci 2019:29–37. Available at: https://doi.org/gr9cb6.

11. Parker HG. Genomic analyses of modern dog breeds. Mamm Genome 2012;23:19–27. Available at: https://doi.org/fz2qt3.

12. Donner J, Freyer J, Davison S, et al. Genetic prevalence and clinical relevance of canine Mendelian disease variants in over one million dogs. Leeb T, ed. PLoS Genet 2023;19:e1010651. Available at: https://doi.org/gr9cb4.

13. Anon. r/DoggyDNA. reddit. Available at: https://www.reddit.com/r/DoggyDNA/. Accessed June 13, 2023.

14. Qin X, Wang Z. NASNet: A Neuron Attention Stage-by-Stage Net for Single Image Deraining. arXiv; 2020. Available at: https://arxiv.org/abs/1912.03151.

15. Parker HG, Dreger DL, Rimbault M, et al. Genomic Analyses Reveal the Influence of Geographic Origin, Migration, and Hybridization on Modern Dog Breed Development. Cell Reports 2017;19:697–708. Available at: https://doi.org/gfpn24.

16. Toro S, Matentzoglu N, Vasilevsky N, et al. monarch-initiative/vertebrate-breed-ontology: v2023-06-01. Zenodo; 2023. Available at: https://doi.org/gsccsz.

17. Anon. Fédération Cynologique Internationale. Available at: https://www.fci.be/en/. Accessed June 13, 2023.

18. Jones S. Best Dog DNA Tests 2023: DNA My Dog vs Wisdom Panel vs Embark vs Orivet vs EasyDNA & More. Canine Journal 2016. Available at: https://www.caninejournal.com/dog-dna-tests-reviews/. Accessed June 13, 2023.

19. Embark Vet. Embark dog DNA test kits. 2023. Available at: https://shop.embarkvet.com/products/embark-dog-dna-test-kit. Accessed June 13, 2023.

20. Deane-Coe PE, Chu ET, Slavney A, et al. Direct-to-consumer DNA testing of 6,000 dogs reveals 98.6-kb duplication associated with blue eyes and heterochromia in Siberian Huskies. Barsh GS, ed. PLoS Genet 2018;14:e1007648. Available at: https://doi.org/gfdqff.

21. Hayward JJ, White ME, Boyle M, et al. Imputation of canine genotype array data using 365 whole-genome sequences improves power of genome-wide association studies. Barsh GS, ed. PLoS Genet 2019;15:e1008003. Available at: https://doi.org/gr9cb3.

22. Sams AJ, Boyko AR. Fine-Scale Resolution of Runs of Homozygosity Reveal Patterns of Inbreeding and Substantial Overlap with Recessive Disease Genotypes in Domestic Dogs. G3 Genes|Genomes|Genetics 2019;9:117–123. Available at: https://doi.org/gr9cb5.

23. Embark Vet. Breed List. 2023. Available at: https://embarkvet.com/resources/dog-breeds/. Accessed June 13, 2023.

24. Wisdom Panel. Wisdom Panel™ Breed Discovery. Available at: https://www.wisdompanel.com/en-us/dog-dna-tests/breed-discovery.

25. Wisdom Panel. The magic of DNA? It lets you meet your dog in an entirely new way. Available at: https://www.wisdompanel.com/en-us/our-science.

26. Orivet. Geno Pet Dog Breed Identification DNA test. Available at: https://www.orivet.com/us. Accessed June 13, 2023.

27. Orivet. Geno Pet Dog Breed Identification Test - DNA Test. Available at: https://www.orivet.com/store/canine-mixed-breed-screen/geno-pet-dog-breed-identification-test. Accessed June 13, 2023.

28. Orivet. Orivet Genetic Pet Care. Available at: https://orivet.com/media/c4ca4238a0b923820dcc509a6f75849b/List%20of%20breeds.pdf.

29. Darwin’s Ark. Darwin’s Ark. Available at: https://darwinsark.org/. Accessed June 13, 2023.

30. Darwin’s Ark. FAQs. Available at: https://darwinsark.org/faqs/. Accessed June 13, 2023.

31. Darwin’s Ark. FAQs. Available at: https://darwinsark.org/faqs/. Accessed June 13, 2023.

32. Omberg L, Salit J, Hackett N, et al. Inferring genome-wide patterns of admixture in Qataris using fifty-five ancestral populations. BMC Genet 2012;13:49. Available at: https://doi.org/gb3dhr.

33. DNA My Dog. See all our tests. DNA My Dog - Fast, Accurate Genetic and Allergy Tests 2022. Available at: https://dnamydog.com/tests/. Accessed June 13, 2023.

34. DNA My Dog. Help Centre and Canine FAQ. Available at: https://dnamydog.com/help/help-centre/. Accessed June 13, 2023.

35. DNA My Dog. Breeds we test. 2022. Available at: https://dnamydog.com/science/breeds-we-test/. Accessed June 13, 2023.

36. accu-metrics. Dog Breed Identification - Identify the Breeds in a Mixed-Breed Dog. accu-metricscom. Available at: https://www.via-pet.com/canine-testing/p/dog-breed-identification. Accessed June 13, 2023.

37. Borwarnginn P, Kusakunniran W, Karnjanapreechakorn S, et al. Knowing Your Dog Breed: Identifying a Dog Breed with Deep Learning. Int J Autom Comput 2020;18:45–54. Available at: https://doi.org/grwfjc.

38. Raduly Z, Sulyok C, Vadaszi Z, et al. Dog Breed Identification Using Deep Learning. In: 2018 IEEE 16th International Symposium on Intelligent Systems and Informatics (SISY). IEEE, 2018. Available at: https://doi.org/grwfjg.

39. Borwarnginn P, Thongkanchorn K, Kanchanapreechakorn S, et al. Breakthrough Conventional Based Approach for Dog Breed Classification Using CNN with Transfer Learning. In: 2019 11th International Conference on Information Technology and Electrical Engineering (ICITEE). IEEE, 2019. Available at: https://doi.org/grwfjf.

40. Anon. PyTorch Image Models. 2023. Available at: https://github.com/huggingface/pytorchimage-models. Accessed June 13, 2023.

41. Wrightman R. Inception ResNet v2. 2021. Available at: https://github.com/huggingface/pytorch-image-models/blob/main/docs/models/inception-resnet-v2.md. Accessed June 27, 2023.

42. Lawson R. Small Sample Confidence Intervals for the Odds Ratio. Communications in Statistics - Simulation and Computation 2004;33:1095–1113. Available at: https://doi.org/fsc7pw.

43. Weber F, Knapp G, Ickstadt K, et al. Zero_cell corrections in random_effects meta_analyses. Res Syn Meth 2020;11:913–919. Available at: https://doi.org/grwfjb.

44. Brzezińska J. The Problem of Zero Cells in the Analysis of Contingency Tables. Zesz Nauk UEK 2015:49–61. Available at: https://doi.org/grwfjh.

45. Meyer D, Zeileis A, Hornik K. The Strucplot Framework: Visualizing Multi-way Contingency Tables with vcd. J Stat Soft 2006;17. Available at: https://doi.org/gfpqfc.

46. Zeileis A, Meyer D, Hornik K. Residual-Based Shadings for Visualizing (Conditional) Independence. Journal of Computational and Graphical Statistics 2007;16:507–525. Available at: https://doi.org/bw646k.

47. Smith MR, Actions-User, Sanselme L, et al. ms609/Ternary: v2.2.0. Zenodo; 2023. Available at: https://doi.org/gsccsx.

48. Khosla A, Jayadevaprakash N, Yao B, Li F. Stanford Dogs dataset for Fine-Grained Visual Categorization. Available at: http://vision.stanford.edu/aditya86/ImageNetDogs/. Accessed June 13, 2023.

49. JoAnn. [New Product] Viaguard DNAffirm Canine DNA Testing Kit. PetMeds® Pet Health Blog 2017. Available at: https://blog.petmeds.com/pet-supplies-products/viaguard-dnaffirm-canine-dna-testing-kit/. Accessed June 13, 2023.

50. Jagannathan V, Drögemüller C, Leeb T, et al. A comprehensive biomedical variant catalogue based on whole genome sequences of 582 dogs and eight wolves. Anim Genet 2019;50:695–704. Available at: https://doi.org/gsccrw.

51. Drever Association of America. Welcome to Drever Association of America. Drever Association of America. Available at: https://dreverusa.org/drever-history. Accessed June 13, 2023.

52. Deutscher Bracken Club e.V. Historie. 2009. Available at: https://web.archive.org/web/20090730055525/ http://www.deutscher-bracken-club.de/pages/die-bracken/historie.php. Accessed June 13, 2023.

53. Ní Leathlobhair M, Perri AR, Irving-Pease EK, et al. The evolutionary history of dogs in the Americas. Science 2018;361:81–85. Available at: https://doi.org/gdvf4h.

54. The McNab Shepherd Registry. The History of the McNab Shepherd from the McNab Shepherd Registry. Available at: https://www.mcnabshepherdregistry.com/dog/history/. Accessed June 13, 2023.

55. Cowley J, News · TD·C. How accurate are dog DNA tests? We unleash the truth | CBC News. CBC 2023. Available at: https://www.cbc.ca/news/business/marketplace-dog-dna-test-1.6763274. Accessed June 13, 2023.

56. Leighton JW, Valverde K, Bernhardt BA. The General Public’s Understanding and Perception of Direct-to-Consumer Genetic Test Results. Public Health Genomics 2011;15:11–21. Available at: https://doi.org/cdbdkb.

57. Leefeldt E, Danise A. Dog Breeds Banned By Home Insurance Companies – Forbes Advisor. 2022. Available at: https://www.forbes.com/advisor/homeowners-insurance/banned-dog-breed-lists/. Accessed June 13, 2023.

58. Donofrio C. Do Insurance Companies Discriminate Against Your Dog? Real Estate News & Insights. 2014. Available at: https://www.realtor.com/advice/buy/insurance-companies-discriminate-against-your-dog/. Accessed June 13, 2023.

59. Deckert T. DNA Testing Your Dog: Why apartment complexes are now making it a requirement. KMEG 2019. Available at: https://siouxlandnews.com/news/local/dna-testing-your-dog-why-apartment-complexes-are-now-making-it-a-requirement. Accessed June 13, 2023.

60. Sanders C. Some Utah landlords want your pet’s DNA. This is the reason why. 2021. The Salt Lake Tribune. Available at: https://www.sltrib.com/news/homeprices/2021/06/18/some-utah-landlords-want/. Accessed June 13, 2023.

61. Zumper. How Do Apartments Verify Dog Breeds? The Zumper Blog 2022. Available at: https://www.zumper.com/blog/how-do-apartments-verify-dog-breeds/. Accessed June 13, 2023.

62. Anon. Home Insurance and Pets. MSPCA-Angell. Available at: https://www.mspca.org/pet_resources/home-insurance-and-pets/. Accessed June 13, 2023.

63. Araj V. A Guide to Homeowners Insurance and Pets. 2021. Available at: https://www.quickenloans.com/learn/homeowners-insurance-and-dogs. Accessed June 13, 2023.

64. George D. How Your Dog Can Impact the Cost of Homeowners Insurance. 2021. Available at: https://www.fool.com/the-ascent/insurance/homeowners/articles/how-your-dog-can-impact-the-cost-of-homeowners-insurance/. Accessed June 13, 2023.

65. Ayers R. Restricted Dog Breeds for Home Insurance. 2023. Available at: https://www.valuepenguin.com/homeowners-insurance-restricted-dog-breeds. Accessed June 13, 2023.

66. Majumder MA, Guerrini CJ, McGuire AL. Direct-to-Consumer Genetic Testing: Value and Risk. Annu Rev Med 2021;72:151–166. Available at: https://doi.org/gh33cv.

67. Ayoubieh H, Blazer K, Christopher D, et al. Direct-to-Consumer Genetic Testing FAQ for Healthcare Professionals. National Human Genome Research Institute. Available at: https://www.genome.gov/For-Health-Professionals/Provider-Genomics-Education-Resources/Healthcare-Provider-Direct-to-Consumer-Genetic-Testing-FAQ. Accessed June 13, 2023.

68. Moses L, Niemi S, Karlsson E. Pet genomics medicine runs wild. Nature 2018;559:470–472. Available at: https://doi.org/gdvnpd.

69. Wall T. DNA tests reveal customized pet food diet for a Mutt. 2016. Available at: https://www.petfoodindustry.com/pet-food-market/article/15462714/dna-tests-reveal-customized-pet-food-diet-for-a-mutt. Accessed June 13, 2023.

70. Embark Vet. Dog Age Test. 2023. Available at: https://shop.embarkvet.com/products/dog-age. Accessed June 13, 2023.

